# Glycogen-Dependent Metabolic Reprogramming Regulates Microglial Activation and Dysfunction in Neurodegenerative Disease

**DOI:** 10.64898/2026.07.13.738226

**Authors:** Hannah McAlister, Heather Merchant, Verity Mitchener, Noemi Gatto, Millie Thackray, Emily Blackburn, Libby Shackleton, Matthew S Gentry, Lorena Arancibia Carcamo, Amy F Lloyd

## Abstract

Microglia are central regulators of neuroinflammation in Alzheimer’s disease (AD), yet how metabolic states modulate function remains unclear. Here we show that microglia from the APP^NL-G-F^ mouse model revealed upregulation of glycolytic enzymes coinciding with onset of microglial activation. Surprisingly, this glycolytic shift occurred alongside reduced expression of glucose transporters, suggesting that extracellular glucose may not be the primary fuel source, implicating glycogenolysis as the potential metabolic driver. Consistent with this, significant microglial glycogen accumulation was noted in late disease, when cells exhibited features of metabolic exhaustion and functional impairment. Pharmacological inhibition of glycogenolysis blunted microglia responses to Abeta aggregates and markedly reduced Abeta uptake, confirming a functional role for glycogen metabolism in shaping microglial states. Together, these findings identify glycogen as a central regulator of microglial metabolic health and function, highlighting glycogen homeostasis as a potential therapeutic target for promoting Abeta clearance and preserving protective microglial functions in AD.

## Introduction

Microglia play a significant role in the initiation and progression of neurodegenerative diseases such as Alzheimer’s disease (AD). AD is pathologically characterised by the presence of amyloid beta (Aβ) plaques and hyperphosphorylated tau neurofibrillary tangles, neuronal and synaptic loss, vascular dysfunction and inflammation (1). As the most common cause of dementia, and a leading cause of death worldwide, treatment options for AD are limited and minimally effective. Furthermore, the primary causes of sporadic AD remain poorly understood.

Genome-wide association studies (GWAS) and genetic profiling have identified many risk loci for late onset AD in genes that are expressed heavily in microglia (2–5). Proteomic profiling of human brain and cerebrospinal fluid (CSF) samples has further identified disease-enriched protein networks associated with microglia and glucose metabolism (6). Furthermore, these protein modules were significantly enriched in AD risk factor loci, highlighting a potential causative role of microglia in AD (6). Understanding the link between microglia and their metabolic states in AD may provide further clues for how microglia contribute to neurodegeneration.

Microglia are both metabolically demanding and flexible; as highly motile cells they are continuously surveying their environment for signs of damage or infection by extending and contracting their processes (7). RNA sequencing of mouse microglia identified expression of genes required for both glycolysis and oxidative phosphorylation (8), confirming the capacity of microglia to use both pathways to generate ATP. Indeed, during homeostasis, microglia use oxidative phosphorylation, however under inflammatory conditions, microglia switch to glycolysis (9–14). This metabolic shift to glycolysis under normoxic conditions is known as the Warburg Effect, initially being discovered as a phenomenon unique to cancer cells to fuel their proliferation (15), however is now appreciated to occur in a range of immune cells upon activation (16). In microglia, this is often accompanied by increases in the expression of glucose transporters (GLUTs) to fuel this metabolic shift (12, 13, 17). Although this switch is thought to occur to meet sudden increases in metabolic demands, it has also been demonstrated that glucose metabolism promotes inflammatory responses via regulation of nuclear factor kappa b (NFκB) transcriptional activity via increased NADH:NAD+ ratio and activation of transcriptional co-repressor CtBP (18). Under low glucose conditions, the NADH:NAD+ ratio decreases, along with reduced transcriptional activity of NFκB and inflammatory gene expression (18). This highlights the importance of metabolic flexibility in driving microglia inflammatory responses.

Although glucose is their primary energy source, microglia are able to switch to glutamine and fatty acid utilisation and metabolism in its absence (19), however the consequences of this on fuelling microglia inflammatory responses are unknown. Furthermore, the long-term consequences of sustained glucose utilisation on microglia health and function also remains poorly understood. Chronic exposure of microglia to Aβ *in vitro* leads to glycolytic and mitochondrial dysfunction, accompanied by reduced inflammatory responses (14). Restoration of glycolysis by mTOR activator interferon gamma (IFN-γ) *in vitro* restored inflammatory responses, and mitigated AD pathology *in vivo* (14), suggesting a benefit of glycolysis-induced inflammatory responses in microglia, however this is met with the risk of metabolic exhaustion and immune tolerance. Indeed, metabolic reprogramming of microglia may contribute to the temporal differences in microglia functions and responses that have been observed in AD over time (20, 21). It is therefore imperative that early and late microglia responses in AD are thoroughly defined to identify novel therapeutic targets to preserve metabolic health and harness the neuroprotective properties of microglia.

As protein expression provides detailed insight into cellular function, we isolated microglia from early and late-stage disease in the APP^NL-G-F^ mouse model of progressive amyloidosis and neurodegeneration, together with age and sex-matched humanised APP controls and characterised them by mass spectrometry. We demonstrate that upregulation of glycolytic pathways accompanies the onset of microglial activation and inflammation but declines as microglia become metabolically dysregulated during later stages of disease. Surprisingly, this increase in glycolytic activity occurred despite a reduction in glucose transporter expression, suggesting that extracellular glucose is unlikely to be the primary fuel source supporting this metabolic shift. Instead, early inflammatory activation was associated with increased expression of enzymes involved in glycogen breakdown. Consistent with this, glycogen accumulation was evident in APP^NL-G-F^ microglia by 12 months of age, when microglia exhibited features of metabolic exhaustion and dysfunction, suggesting impaired glycogen mobilisation. Finally, pharmacological inhibition of glycogen phosphorylase, the rate-limiting enzyme of glycogenolysis, attenuated the induction of key amyloid-responsive proteins, and markedly reduced Aβ uptake. Collectively, these findings identify glycogen mobilisation as a previously unrecognised regulator of microglial metabolic fitness and the microglial response to amyloid pathology. Targeting glycogen metabolism may therefore represent a novel therapeutic strategy to preserve microglial function and promote protective immune responses in neurodegenerative disease.

## Results

### Analysis of APP^NL-G-F^ Microglia Shows Striking Proteomic Reprogramming with Age and Disease Progression

Microglia were isolated from the APP^NL-G-F^ mouse model along with sex and aged-matched controls from mice carrying the non-pathogenic humanised APP (hAPP) gene, providing a more robust control and measure of pathogenic APP compared to wild-type mice. Detailed descriptions of each sample and group can be found in Supplemental File 1. No significant differences in isolated cell numbers were found between ages, genotype or sex (Supplemental File 1). Microglia were isolated based on documented pathological events (22), enabling detailed analysis of microglia from prodromal (1-month), onset of gliosis (3-month), significant plaque development and cognitive impairment (6-month) and late disease stages (12-month) (Figure 1a). A total of 8843 proteins were identified across all samples. Microglia protein mass was between 30 and 40 picograms per cell in all samples (Figure 1b).

**Figure 1.**
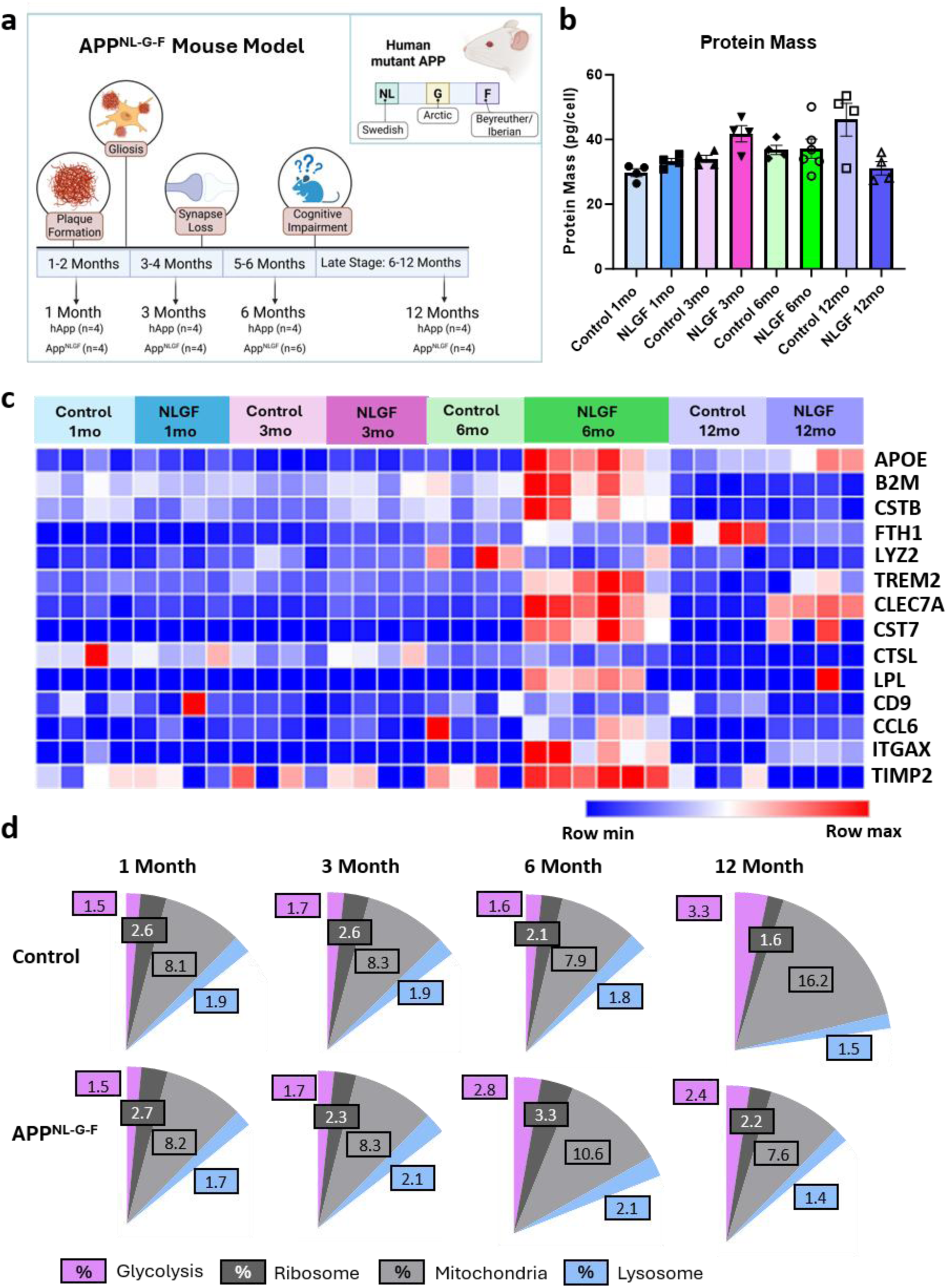
Proteomic analysis of APP^NL-G-F^ microglia highlights dynamic proteome reprogramming throughout disease progression. **a)** Schematic of the APP^NL-G-F^ mouse model and chosen time points for microglia analysis. Created using BioRender.com. **b)** Protein mass of all microglia samples from APP^NL-G-F^ mice and time-matched controls ± S.E.M. N=4-6 mice per time point**. c)** Heatmap of disease-associated microglia (DAM) protein expression across all time points. Red = upregulated, blue = downregulated. **d)** Pie charts representing proteins associated with glycolysis, ribosomes, mitochondria and lysosomes as a proportion of total proteome (%) in control and APP^NL-G-F^ microglia across all time points.

Disease-associated microglia (DAM; (23)) protein expression was compared across all samples, highlighting dynamic changes to this protein network with disease progression (Figure 1c). Most DAM-associated proteins, including APOE, B2M, CSTB, TREM2, CLEC7A, CST7, LPL, ITGAX and TIMP2 were strongly upregulated in 6-month APP^NL-G-F^ mice relative to all other groups. Interestingly, by 12 months, only CLEC7A and to a lesser extent, APOE, remained consistently upregulated in APP^NL-G-F^ microglia, with all other DAM proteins decreasing in expression (Figure 1c). Changes in proteins associated with metabolic and core subcellular components were also observed, noted in the proportion of the proteome dedicated to glycolysis, ribosome, mitochondria, and lysosome machinery (Figure 1d). As well as a strong DAM signature, 6-month APP^NL-G-F^ microglia showed increases in proportions of the proteome dedicated to glycolysis, ribosome and mitochondria machinery compared to 6-month controls and earlier APP^NL-G-F^ and control time points. By 12 months, these proportions were reduced. Interestingly, 12-month hAPP control microglia show noted increases in glycolysis and mitochondrial protein expression, suggesting that age-related proteomic reprogramming is also evident, and therefore downregulation of these components in diseased microglia is even more remarkable.

Together, these findings demonstrate a temporally dynamic microglial proteomic response in APP^NL-G-F^ mice, characterised by a pronounced DAM and metabolic activation at mid-stage disease that subsequently attenuates in late-stage pathology, alongside distinct age-associated proteomic adaptations.

### Proteomic Analysis Reveals Differences in Metabolic and Inflammatory Microglia Signatures with Disease Progression

Differential expression analysis of proteins expressed in APP^NL-G-F^ microglia compared to age-matched hAPP controls revealed no significant changes in protein expression at 1 and 3 months (P-Adj>0.05; Supplemental Figure 1a, b). However, by 6 months, striking proteomic changes were observed, such that 37% of all proteins were significantly enriched in APP^NL-G-F^ microglia compared to control, with 2% significantly downregulated at 61% of proteins unchanged (Figure 2a). This included rate-limiting glycolytic enzymes which were shown to be strongly upregulated in APP^NL-G-F^ microglia at this time (Figure 2b, Supplemental Figure 2). DAM proteins ITGAX and CLEC7A were among the most significantly enriched in 6-month APP^NL-G-F^ microglia, as well as proteins that participate in glucose and fatty acid metabolism, TP53-inducible glycolysis and apoptosis regulator (TIGAR) and Acetyl-CoA Carboxylase Alpha (ACACA), respectively (Figure 2c). Interestingly, despite noted increases in glycolytic enzymes and associated proteins, glucose transporters SLC2A1 (GLUT1) and SLC2A3 (GLUT3) were significantly downregulated (Figure 2c, Supplemental File 1), which was surprising given that increases in glycolysis are usually associated with increases in glucose transporter expression (24). Other glucose transporters such as SLC2A2 (GLUT2) and SLC2A4 (GLUT4) were not detected. Gene Ontology (GO) term analysis revealed upregulation of proteins associated with inflammatory responses in APP^NL-G-F^ microglia at 6 months compared to controls (Figure 2d) and downregulation of glycosylation and glycolipid metabolism (Figure 2e).

**Figure 2.**
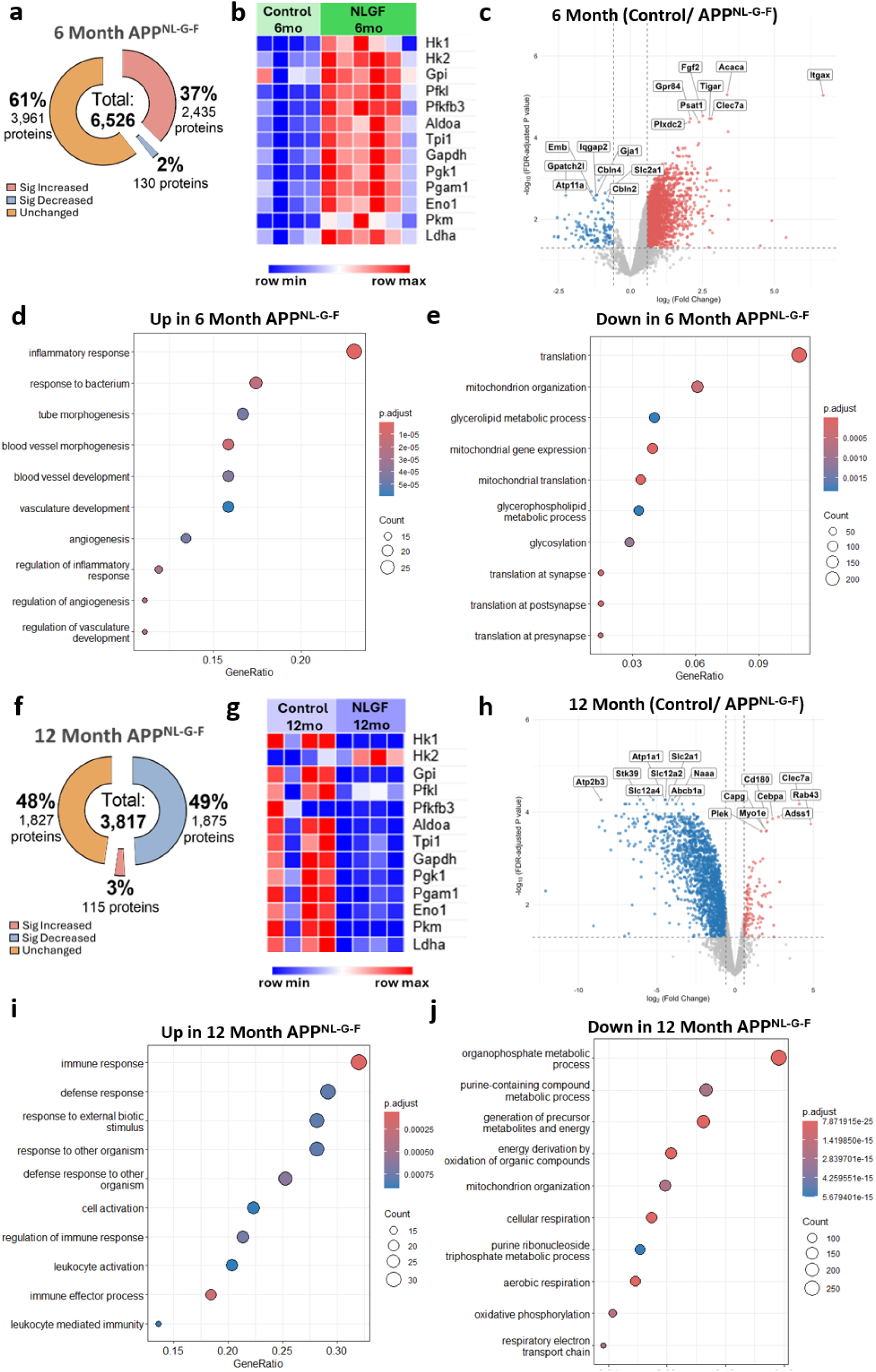
Late-stage APP^NL-G-F^ microglia are associated with dysregulated metabolic programming and reduction in amyloid response protein expression. **a)** Pie chart highlighting the total number of proteins that are significantly enriched (red), significantly downregulated (blue) or unchanged (orange) in 6-month APP^NL-G-F^ microglia compared to time-matched control. **b)** Heatmap of glycolysis enzyme expression in 6-month APP^NL-G-F^ microglia and time-matched control. Red = increased expression, blue = decreased expression. **c)** Volcano plot of differentially expressed proteins between 6-month APP^NL-G-F^ microglia and time-matched control. Red dots = significantly upregulated proteins in APP^NL-G-F^ microglia (FC > 1.5, FDR-adjusted P-value <0.05), blue dots = significantly downregulated proteins in APP^NL-G-F^ microglia (FC > 1.5, FDR-adjusted P-value <0.05). **d)** Significantly enriched GO terms in 6-month APP^NL-G-F^ microglia. **e)** Significantly downregulated GO terms in 6-month APP^NL-G-F^ microglia. **f)** Pie chart highlighting the total number of proteins that are significantly enriched (red), significantly downregulated (blue) or unchanged (orange) in 12-month APP^NL-G-F^ microglia compared to time-matched control. **g)** Heatmap of glycolysis enzyme expression in 12-month APP^NL-G-F^ microglia and time-matched control. Red = increased expression, blue = decreased expression. **h)** Volcano plot of differentially expressed proteins between 12-month APP^NL-G-F^ microglia and time-matched control. Red dots = significantly upregulated proteins in APP^NL-G-F^ microglia (FC > 1.5, FDR-adjusted P-value <0.05), blue dots = significantly downregulated proteins in APP^NL-G-F^ microglia (FC > 1.5, FDR-adjusted P-value <0.05). **i)** Significantly GO terms in 12-month APP^NL-G-F^ microglia. **j)** Significantly downregulated GO terms in 12-month APP^NL-G-F^ microglia.

In contrast, at 12 months, 49% of proteins were significantly downregulated in APP^NL-G-F^ compared to control, with only 3% of proteins significantly enriched and 48% unchanged (Figure 2f). Compared to control, most glycolytic enzymes (except for HK2) were downregulated in 12-month APP^NL-G-F^ microglia (Figure 2g, Supplemental Figure 2) and global differential analysis revealed continual downregulation of SLC2A1 and SLC2A3 along with solute transporters SLC12A2 and SLC12A4 (Figure 2h, Supplemental File 1). DAM-associated protein CLEC7A remained significantly upregulated in 12-month APP^NL-G-F^ microglia (Figure 2h) however all other DAM proteins were unchanged compared to control. GO term analysis indicated that immune and inflammatory responses were upregulated (Figure 2i), whereas proteins associated with cellular respiration and metabolism were downregulated (Figure 2j). These data suggest that although APP^NL-G-F^ microglia at 12 months are still inflammatory, this response is associated with metabolic dysregulation.

### Dysregulated Microglia Responses and Features of Senescence in Later Stages of Disease

Differential expression analysis comparing 6- and 12-month APP^NL-G-F^ microglia was performed to define the phenotypic and functional changes occurring during disease progression. Although 12-month APP^NL-G-F^ microglia show significant enrichment of CLEC7A compared to time-matched controls, expression is still significantly lower than 6-month APP^NL-G-F^ microglia (Supplemental Figure 3a).

Repeated exposure of microglia to Aβ aggregates induces immune tolerance and metabolic exhaustion *in vitro* (14). Reduction of DAM proteins and lysosomal machinery suggest a similar metabolically exhausted phenotype is present at 12 months in the APP^NL-G-F^ model. To look at this further we identified some key proteins associated with metabolic fitness. TREM2, important for both DAM phenotype engagement (23) and metabolic fitness (24) is significantly upregulated in 6-month APP^NL-G-F^ microglia, then decreases by 12 months (Supplemental Figure 3b). Sequestosome 1 (SQSTM1; p62), a selective autophagy receptor involved in clearing protein aggregates and processing glycogen (25) is upregulated at 6 months in APP^NL-G-F^ microglia but not at 12 months (Supplemental Figure 3c), suggestive of reduced autophagy capacity. As nutrient stress decreased autophagy and exhaustion is linked to senescence, we looked for evidence of this and identified upregulation of H2AX, a DNA damage response protein, in 12-month APP^NL-G-F^ microglia (Supplemental Figure 3d). The phosphorylated (γ) form of H2AX is associated with persistent DNA double strand breaks and cellular senescence. Therefore, to confirm expression, immunostaining of γ-H2AX was conducted and found to be significantly increased in microglia at 12 months compared to 6 months (Supplemental Figure 3e, f). This data therefore highlights the link between metabolic fitness and the capacity to engage with and clear amyloid, with microglia with signs of DNA damage more prevalent in later stages of disease.

### Dysregulated Glycogen Metabolism is a Feature of Late-Stage Disease in APP^NL-G-F^ Microglia

The observed downregulation of glucose transporters, together with the transient induction and subsequent decline of glycolytic enzymes during disease progression, led us to hypothesise that microglia utilise intracellular glycogen stores, rather than extracellular glucose, to support their metabolic demands. To test the contribution of extracellular glucose to microglial function, primary mouse microglia were cultured in glucose-deprived medium for 24 hours. Surprisingly, glucose deprivation induced minimal changes to the basal microglial proteome compared with cells maintained in complete medium (17 mM glucose; Supplemental Figure 4a). Moreover, the proteomic response to lipopolysaccharide (LPS) stimulation was preserved under glucose-deprived conditions, with no significant differences detected between LPS-treated microglia cultured in the presence or absence of glucose (Supplemental Figure 4b). Together, these findings indicate that acute inflammatory activation of microglia can occur largely independently of extracellular glucose availability, supporting the hypothesis that intracellular energy reserves may sustain microglial metabolic responses.

To investigate this further, we examined the expression of key enzymes involved in glycogen metabolism, encompassing both glycogenolysis and glycogenesis (Figure 3a). This revealed marked changes in glycogen phosphorylase (PYG) and phosphoglucomutase (PGM1), two enzymes essential for mobilising glycogen-derived glucose (Figure 3b). Both proteins were increased by approximately twofold in 6-month APP^NL-G-F^ microglia relative to controls but were reduced by approximately fourfold in 12-month APP^NL-G-F^ microglia (Figure 3b). Expression of glycogen synthase (GYS1), the rate-limiting enzyme in glycogen synthesis and a marker of cellular glycogen homeostasis, was also markedly decreased at 12 months. Other enzymes involved in the formation of glycogen, UDGP Pyrophosphorylase (UDP2), Glycogenin (GYG1) and Glycogen Branching Enzyme (GBE1) were detected, with UGP2 and GBE1 showing the lowest expression in 12-month APP^NL-G-F^ microglia (Supplemental Figure 5a). GBE1 in particular was most highly expressed in 6-month APP^NL-G-F^ microglia (Supplemental Figure 5a). Glycogen Debranching Enzyme (GDE/ AGL) was also least expressed in 12-month APP^NL-G-F^ microglia (Supplemental Figure 5b), and all detected subunits of phosphorylase kinase, the group of enzymes vital for PYG activation, were lowly expressed or undetected in 12-month APP^NL-G-F^ microglia compared to other time points and controls (Supplemental Figure 5c), suggesting that glycogen breakdown is impaired. Together, these findings indicate profound dysregulation of glycogen metabolism during disease progression and suggest that impaired glycogen mobilisation or turnover contributes to the metabolically exhausted microglial phenotype.

**Figure 3.**
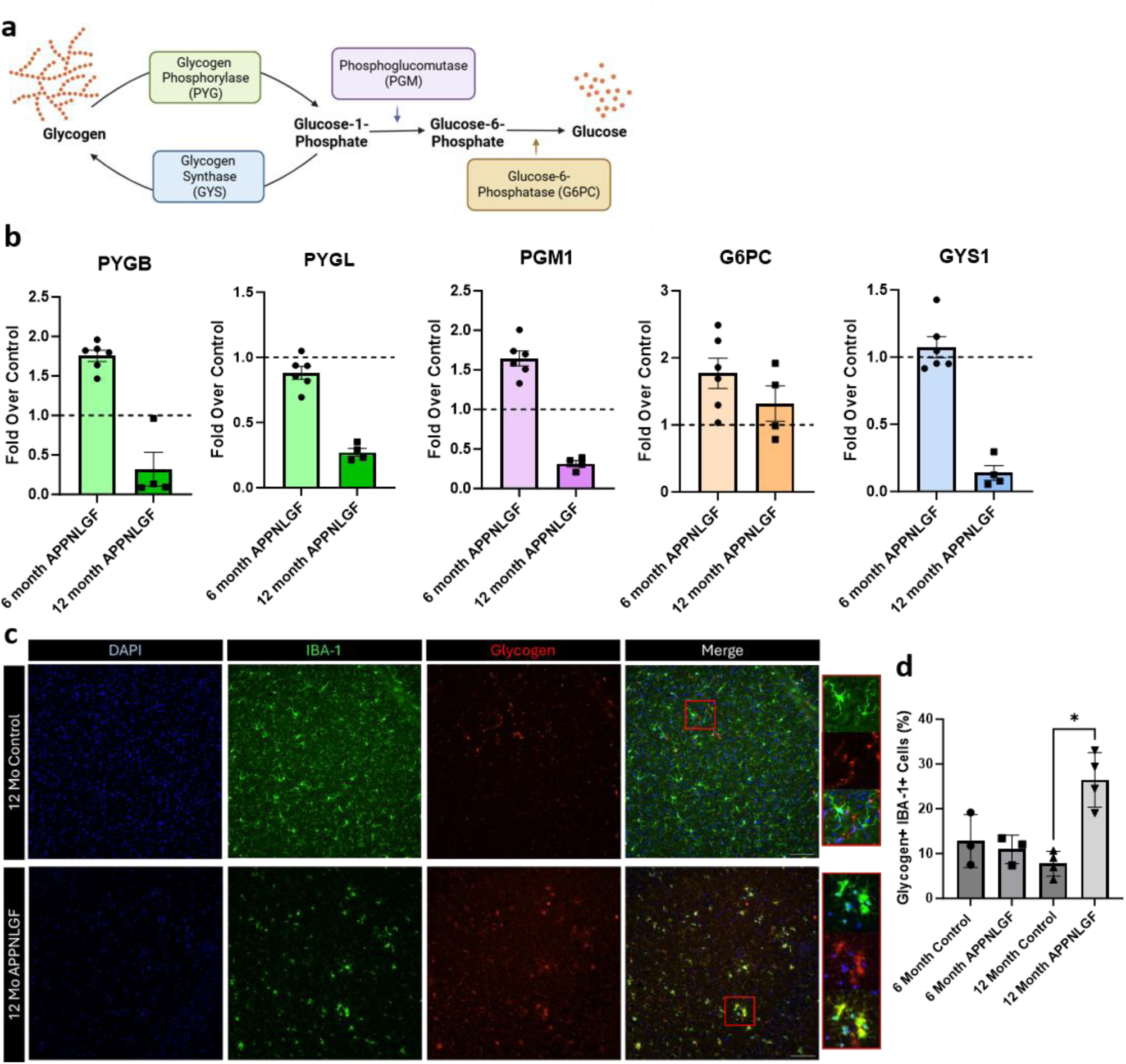
Dysregulated glycogen metabolism is associated with glycogen accumulation in late-stage APP^NL-G-F^ microglia. **a)** Schematic of enzymes associated with glycogenolysis (PYG, PGM, G6PC) and glycogenesis (GYS). Created using BioRender.com. **b)** Relative protein expression of enzymes associated with glycogenolysis (PYGB (brain form), PYGL (liver form), PGM, G6PC) and glycogenesis (GYS) in 6- and 12-month APP^NL-G-F^ microglia compared to age-matched controls (dotted lines) ± S.E.M. N=4-6 mice per time point. **c)** Representative images from immune-stained cortical brain sections from 12-month APP^NL-G-F^ and control mice. Blue = DAPI (nuclei), Green = IBA-1 (microglia), Red = glycogen. Scale bar = 100µm. Right insert = magnified section highlighting glycogen accumulation observed in APP^NL-G-F^ IBA-1+ microglia but not in time-matched control. **d)** Quantification of glycogen+ IBA-1+ microglia from 6- (N=3) and 12-month (N=4) APP^NL-G-F^ brains and time-matched controls ± S.E.M.

To validate the observed alterations in glycogen metabolism, we quantified glycogen content in microglia within APP^NL-G-F^ brain tissue. Using a previously validated anti-glycogen antibody (26) we detected significantly increased numbers of glycogen+ microglia in the APP^NL-G-F^ mice at 12 months (Figure 3c, d). These findings are consistent with our proteomic analysis and indicate that glycogen accumulates in microglia during the later stages of disease, supporting the hypothesis that impaired glycogen mobilisation contributes to microglial dysfunction.

### Inhibition of Glycogen Breakdown Attenuates Phagocytosis of Aβ in Microglia *In Vitro*

As glycogen-positive puncta were detected in non-microglia cells, likely reflecting the glycogen reserves of neurons and astrocytes (25) it was important to determine whether the glycogen accumulation observed in late-stage disease microglia resulted from impaired glycogen metabolism or a consequence of phagocytosing glycogen-rich cells. Therefore, we used *in vitro* mouse microglia monocultures to directly measure glycogen-dependent microglia functions in response to Aβ aggregate exposure (Figure 4a).

**Figure 4.**
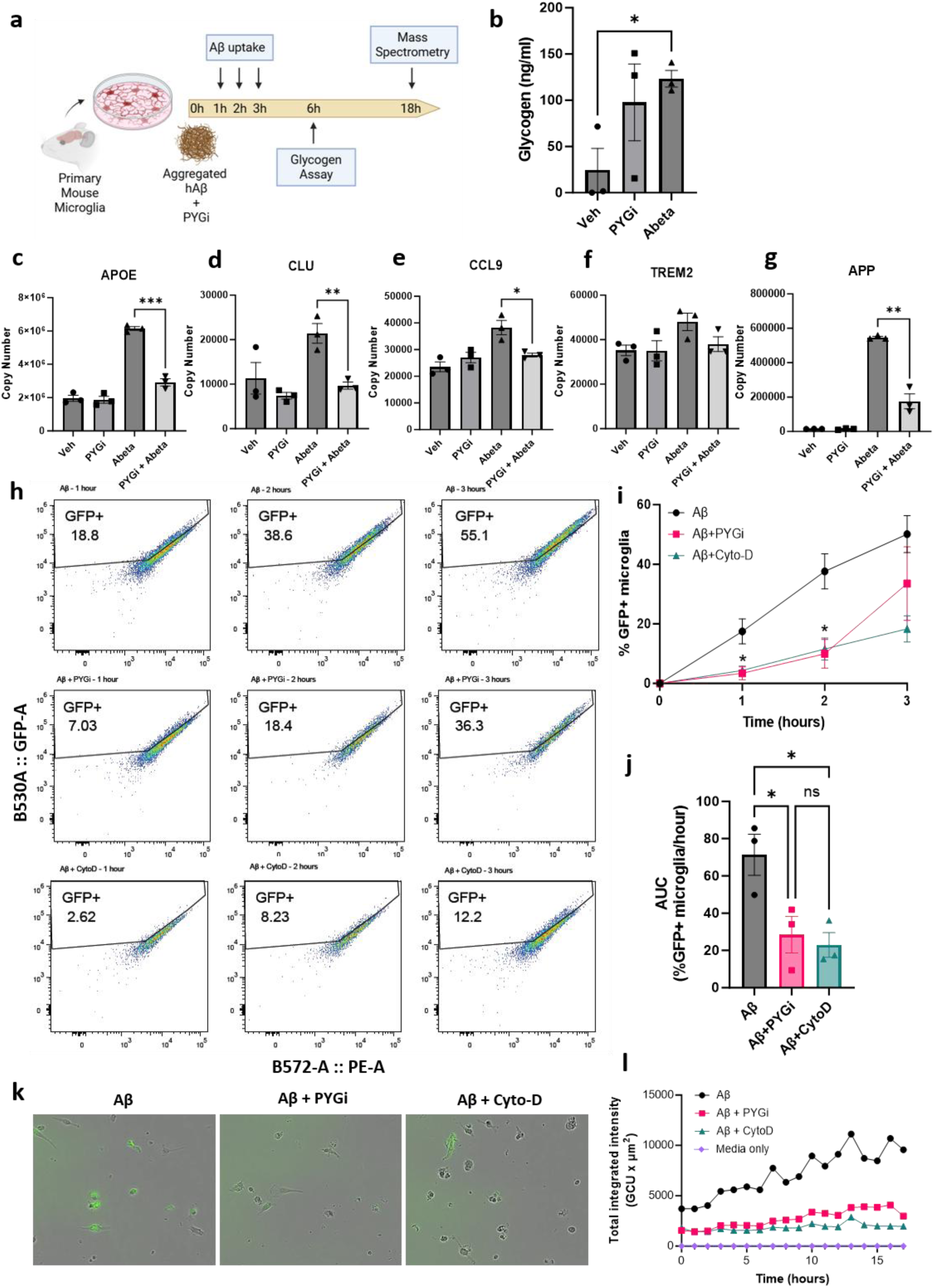
Preventing glycogen breakdown in microglia inhibits DAM phenotype and inhibits Aβ uptake. **a)** Schematic of *in vitro* mouse microglia studies to determine importance of glycogen mobilisation on Aβ responses. Created using BioRender.com. **b)** Quantification of intracellular glycogen concentrations in microglia under 6 hours of vehicle (DMSO) control, inhibition of PYGi and exposure to Aβ ± S.E.M. N=3. *p<0.05 (Students T-Test). **c)** Proteomic analysis of APOE copy numbers in vehicle (DMSO), PYGi, Aβ and Aβ + PYGi ± S.E.M. N=3. *** p<0.001 (Students T-Test). **d)** Proteomic analysis of Clusterin (CLU) copy numbers in vehicle (DMSO), PYGi, Aβ and Aβ + PYGi ± S.E.M. N=3. ** p<0.01 (Students T-Test). **e)** Proteomic analysis of CCL9 copy numbers in vehicle (DMSO), PYGi, Aβ and Aβ + PYGi ± S.E.M. N=3. * p<0.05 (Students T-Test). **f)** Proteomic analysis of TREM2 copy numbers in vehicle (DMSO), PYGi, Aβ and Aβ + PYGi ± S.E.M. N=3. **g)** Proteomic analysis of amyloid precursor protein (APP) copy numbers in vehicle (DMSO), PYGi, Aβ and Aβ + PYGi ± S.E.M. N=3. ** p<0.01 (Students T-Test). **h)** Representative flow cytometry plots of primary mouse microglia after 1, 2 and 3 hours of exposure to GFP-Aβ aggregates. Top row: Aβ alone; middle row: Aβ + glycogen phosphorylase inhibitor (PYGi); bottom row: Aβ + phagocytosis inhibitor Cytochalasin-D (Cyto-D). **i)** Quantification of flow cytometry analysis measuring percentage of GFP+ microglia over time ± S.E.M. N=3. * p<0.05 (Students T-Test; Aβ v Aβ+PYGi). **j)** Quantification of the area under the curve (AUC) of flow cytometry time course data ± S.E.M. *p<0.05 (One-way ANOVA, Tukey’s Multiple Comparisons post-hoc test). N=3. **k)** Time-matched frames from Incucyte real-time analysis of GFP-Aβ uptake in primary mouse microglia. Left panel = Aβ only; middle panel = Aβ + PYGi; right panel = Aβ + Cyto-D. **l)** Quantified rate of uptake (GFP intensity of cells over time) of GFP-Aβ in primary mouse microglia from Incucyte image analysis. N=1.

Firstly, quantification of intracellular glycogen revealed accumulation in microglia 6 hours after Aβ exposure (Figure 4b) at similar levels to microglia treated with the PYG inhibitor CP-91149 (27) (PYGi), implicating glycogen breakdown as a dynamic metabolic process that may be disrupted by Aβ exposure. Furthermore, mass spectrometry analysis of microglia proteomes revealed that although most of the proteome was unchanged between Aβ treated and Aβ + PYGi microglia, there were significant differences in expression of key amyloid response proteins. As expected, Aβ exposure induced expression of DAM-associated proteins APOE (Figure 4c),Clusterin (CLU; Figure 4d), CCL9 (Figure 4e) and TREM2 (Figure 4f), however this was diminished by co-treatment with PYGi (significantly except for TREM2). Furthermore, copies of APP, indicative of Aβ, were also significantly decreased in Aβ + PYGi treated microglia compared to Aβ alone (Figure 4g). This data therefore suggests that glycogen metabolism may be vital in regulating microglia responses to Aβ.

Finally, to confirm the importance of glycogen metabolism on Aβ responses, we quantified rates of Aβ uptake in *in vitro* mouse microglia using GFP-tagged aggregated Aβ and flow cytometry (gating strategy found in Supplemental Figure 6a). Flow cytometry analysis revealed that the proportion of GFP+ microglia increased steadily within the first 3 hours of Aβ exposure, with phagocytosis inhibitor Cytochalasin-D significantly attenuating uptake (Figure 4h - j), highlighting phagocytosis as a major method of Aβ clearance in microglia. Similarly, PYGi also caused a significant reduction in Aβ uptake, confirming an important role of glycogen metabolism on Aβ clearance (Figure 4h - j). This striking inhibition was also observed by live imaging (Figure 4k, l, Supplemental Video 1). Inhibitors alone did not cause any autofluorescence that could account for the increase in GFP+ microglia (Supplemental Figure 6b). PYGi also did not alter phagocytosis rates of pHrodo red E. coli bioparticle uptake in primary microglia (Supplemental Figure 6c, d), suggesting that PYG inhibition does not broadly impact phagocytosis, therefore specifically highlighting Aβ clearance in microglia as a glycogen-sensitive mechanism. Furthermore, PYGi did not directly affect the stability of GFP-Aβ as it did not inhibit the increase in GFP+ BV2 cells over time when given the same GFP-Aβ aggregates (Supplemental Figure 7a, b). Interestingly, 2-deoxyglucose (2-DG), a competitive inhibitor of glycolysis, did reduce Aβ uptake in BV2 cells to similar levels as Cytochalasin-D (Supplemental Figure 7a, b). This data also highlights a fundamental difference in metabolic demand between primary microglia and BV2 cells that supports the proteomic differences previously investigated by our lab (28).

Altogether, this data implicates glycogen metabolism as a vital regulator of Aβ responses and clearance mechanisms in microglia, with dysregulated glycogen metabolism a contributor of microglial metabolic exhaustion, failed clearance, and Aβ plaque accumulation.

## Discussion

In this study, we define stage-specific metabolic reprogramming of microglia in the APP^NL-G-F^ mouse model and identify glycogen metabolism as a central regulator of Aβ-associated microglial activation and functional integrity. Using proteomic profiling across early and late disease stages, we demonstrate that upregulation of glycolysis enzymes coincides with inflammatory activation in early pathology but declines as microglia transition toward a metabolically exhausted and dysfunctional phenotype. Importantly, this glycolytic shift occurred independently of increased glucose transporter expression, instead associating with enhanced glycogenolysis, revealing glycogen as a previously underappreciated endogenous fuel source supporting microglial function.

Microglial metabolic switching toward glycolysis under inflammatory conditions is well established (14, 18, 29), however, the source of substrate supporting this transition has remained unclear. Other AD associated proteome datasets from mouse (30) and human (31) have identified significant upregulation of proteins associated with glycolysis in microglia. Our data suggests that intracellular glycogen, rather than extracellular glucose, sustains early inflammatory and functional responses. The observed downregulation of glucose transporters alongside increased glycolytic enzyme abundance challenges the prevailing assumption that enhanced glucose uptake is required for microglial activation. Indeed, increased glucose influx is important for glycolysis and immune functions in adaptive immune cells, with glucose deprivation affecting cellular functions within hours (32, 33). In microglia, proteomes are unaffected after 24 hours of glucose deprivation, with no differences in the engagement of LPS-induced inflammatory phenotypes, further supporting glycogen metabolism as an essential mediator of microglia metabolic fitness and function.

These findings provide a potential mechanistic explanation for the temporal evolution of microglial phenotypes observed in Alzheimer’s disease. Early activation may confer protective functions, including plaque containment and debris clearance, supported by glycogen-fuelled glycolysis and/or glycogen-mediated signalling, which becomes dysregulated as disease progresses. Moreover, dark microglia, defined by their distinct morphology and localisation to plaques and synapses (34) possess glycogen granules as observed by electron microscopy (35), consistent with late-stage glycogen-filled plaque-associated microglia in our study. Failure of glycogen breakdown may therefore be a key driver of microglial metabolic exhaustion, contributing to the impaired and dysregulated states characteristic of late disease stage.

Functionally, inhibition of glycogen metabolism by inhibition of PYG specifically altered expression of Aβ-associated proteins, most notably APOE. This, along with its ability to inhibit Aβ uptake in microglia, suggests that glycogen may have some regulatory impact on APOE which in turn reduces APOE-mediated uptake of Aβ. APOE has been reported to impact brain metabolic homeostasis in an isoform-dependent manner (36, 37). For example, APOE can modulate insulin signalling and glycolysis in astrocytes (37) yet its role in microglia is not as well defined. Furthermore, a direct link between glycogen metabolism and APOE has not previously been reported. Similarly, Clusterin, another Aβ-binding molecular chaperone was also shown to be modulated by glycogen metabolism. Understanding how glycogen mobilisation and glucose availability regulates Aβ-associated protein expression and Aβ uptake will be a vital next step to understand how to therapeutically restore Aβ clearance in metabolically exhausted microglia, and if this impacts broader protein aggregate clearance in other neurodegenerative diseases. Our data suggests that promoting microglial glycogen metabolism could sustain beneficial responses during early neurodegeneration, while preventing exhaustion. Targeting metabolic pathways in microglia offers a strategy distinct from broadly suppressing inflammation, instead aiming to optimise functional capacity. Indeed, restoring glycolysis with IFN-γ treatment of *in vitro* microglia restores metabolic health and functional responses to Aβ (14). How these experiments impacted glycogen are unclear, however one hypothesis could be that interferon signalling promotes the mobilisation of glycogen to fuel metabolic demands. Furthermore, further studies are required to determine whether modulation of glycogen metabolism *in vivo* can alter Aβ burden, neuronal integrity, or cognitive outcomes.

Together, our findings provide novel insight into the metabolic landscape of microglia in neurodegeneration and highlight glycogen homeostasis as a promising target for preserving protective, Aβ-clearing microglial functions in Alzheimer’s disease.

## Methods and Materials

### Mice

APP^NL-G-F^ and APP^Hu^ (hAPP) mice were used in this study in accordance with the UK Animal (Scientific Procedures) Act, 1986. Experimental procures were approved by the UK Home Office and ethical approval was granted through review by a local advisory committee (PP4652483). For *in vitro* studies wild-type C57-BL6 mice (Jackson Labs) aged 3-4 months were used. For collection of tissue, mice were anaesthetised with a fatal dose of 70% pentobarbital in saline (60mg/kg) and monitored until loss of pedal reflex. They were then trans-cardinally perfused with ice-cold 1X DPBS supplemented with 5U of heparin and the brain was dissected. Further details of mice used for mass spectrometry and immunohistochemistry are found in Supplemental File 1. Hemispheres kept for histology were immersion fixed in 4% PFA overnight at 4°C. Brains were then cryoprotected in 30% sucrose solution overnight at 4°C before being embedded in OCT and stored at -80°C until sectioning.

### Isolation of APP^NL-G-F^ and hAPP Control Mouse Microglia

Following perfusion, the cerebellum and olfactory bulbs were removed, and the brain was placed in cold FACS buffer (1 × DPBS, 2% BSA and 2 mM EDTA). Brains were mechanically and enzymatically dissociated using Miltenyi Neural Tissue Dissociation Kit P as per manufacturer’s instructions (Miltenyi, 130-092-628). Samples were passed through a 70-μm strainer (Greiner, 542070), washed in 15 ml of ice-cold FACS buffer and spun at 300g for 15 minutes. Pellets were resuspended in 30% isotonic Percoll (GE Healthcare, 17-5445-02) and centrifuged at 300g for 15 minutes to remove myelin. The myelin layer was aspirated and discarded. Pellets were resuspended in cold FACS buffer containing FcR blocking reagent (1:10, Miltenyi, 130-092-575) for 10 min. Next, cells were incubated with the following antibodies: PE-Pan-CD11b (1:50, Miltenyi, 130-113-806), BV421-mCD45 (1:500, BD Biosciences, 563890) and viability dye (1:2000, eFluor 780, Thermo Fisher, 65-0865-14), in cold FACS buffer for 30 minutes. After incubation, cells were washed, and the pellet was resuspended in 1mL FACS buffer, passed through a 35μm strainer, and sorted on a MACSQuant Tyto. The cell suspension was loaded into the input chamber of a MACSQuant® Tyto® HS Cartridge (Miltenyi, 130-121-549) and microglia were sorted based on CD11b⁺CD45^low expression. All steps were conducted at 4°C.

### Isolation and Culture of *In Vitro* Mouse Microglia

Mice were culled by CO_2_ and femoral cut, and brains were collected in ice-cold HBSS. Tissue was processed as described above. For microglia isolation for *in vitro* cultures, the single cell suspension was then incubated on ice with CD11b Microbeads (Miltenyi, 130-097-142) for 15 minutes before being washed with MACS buffer (2% BSA, 1% EDTA in PBS). Cells were then run through pre-rinsed (with MACS buffer) LS columns (Miltenyi, 130-042-401) attached to a QuadroMACS™ Separator magnet (Miltenyi, 130-091-051). After washing with 12ml MACS buffer, columns were removed from the magnet and cells retained (microglia) were flushed in 5ml MACS buffer, before a final spin at 400g for 5 minutes. All steps were conducted at 4°C.Microglia were resuspended in warm Dulbecco’s Modified Eagle’s Medium/Nutrient Mixture F-12 (DMEM/F-12) supplemented with 100 U/ml penicillin and 100 mg/ml streptomycin (Sigma, P4333), 10% heat-inactivated foetal bovine serum (FBS; Thermo Fisher, A5256701), 500 ng/ml rhTGFb-1 (Miltenyi, 130-095-066), 10 ng/ml mCSF1 (RCD Systems, 416-ML-050/CF). Microglia were counted using a haemocytometer and plated out onto 24-well plates (Corning) at a density of 125,000 cells per well for flow cytometry, or onto 48-well plates (Corning) at a density of 50,000 cells per well for Incucyte analysis and incubated at 37°C at 5% CO_2_ with high humidity.

### Culture of BV2 cells

BV2 cells were cultured in RPMI 1640 (Gibco, 11875093) with 10% foetal bovine serum (FBS) at 37°C with 5% CO2, with passaging every 2–3 days with trypsin/EDTA (Gibco, 25200056). For flow cytometry experiments, cells were seeded at 100,000 cells per well of a 24 well plate and left to adhere and proliferate overnight before experimentation.

### Preparation of GFP^+^ Aβ_42_

Alexa Fluor 488 labelled Aβ_42_ (AnaSpec, AS-60479-01) was reconstituted in Hexafluoroisopropanol (HFIP; Sigma, 105228) solvent, dried down for 1 hour in a SpeedVac and stored at -20°C. To produce fibrillar Aβ, 2µl of DMSO was added to 45µg Aβ_42_, sample vortexed and centrifuged briefly before sonicating for 10 minutes. 98µl of HCl was added before incubating at 37°C for 24 hours to induce fibrillar aggregation.

### Aβ uptake experiments (flow cytometry)

Cultures were treated with Aβ alone (0.2µM) or Aβ ± 10µM CP-91149 (glycogen phosphorylase inhibitor (PYGi) SelleckChem, S2717) or 10µM Cytochalasin D (Merck, C8273). Cells treated with the inhibitors received a 30-minute pre-treatment before the addition of Aβ ± inhibitors for 1, 2, or 3 hours. Following treatment, media was collected in FACS tubes before washing cells with 1X HBSS (Gibco, 14025092). Trypsin-EDTA (Gibco, 25300054) was added and cells incubated at 37°C for 5 minutes. 1 ml FACS buffer (1X HBSS, 1% EDTA, and 2% BSA) was added to neutralise the trypsin and cells were transferred to FACS tubes. Samples were centrifuged at 300g for 5 minutes, supernatant discarded and cells resuspended in 300µl FACS buffer ± DAPI. Samples were analysed using a NovoCyte flow cytometer (Agilent) and gating analysis was performed using FlowJo (version 10.10). Live cells were gated using DAPI negative against SSC-A, doublets excluded using FSC-A and width, followed by % GFP+ cells gated using GFP-A and PE-A for autofluorescence analysis.

### Real-time Aβ_42_ uptake assay (Incucyte)

50,000 primary adult microglia were seeded per well of a 48-well plate. Following 24 hours in culture, cells were treated with Aβ alone or Aβ ± PYGi (10µM) or cytochalasin D (10µM) and imaged every hour for 24 hours using an Incucyte S3 Live Cell Analysis Instrument (Sartorius). An image of untreated cells was taken before the addition of treatments. Videos were created and rate of uptake was measured using Incucyte Analysis Software (v2024B, Sartorius).

### Immunohistochemistry

OCT-embedded brains were coronally sectioned to 10µm thick using a cryostat, with 5 sections per slide taken around -3.08mm for sections to include the hippocampus (Allen Mouse Brain Atlas) and stored at -80°C until use. Sections were taken out of the -80°C freezer and left to get to room temperature (RT) for 15 minutes. Sections were then given 150µl of blocking buffer (0.2% Triton-X-100 (Sigma, X100), 5% BSA (Sigma, A7979) in PBS) for 1 hour at RT to permeabilise the slides and block non-specific binding. Primary antibodies were then diluted in blocking buffer and added to slides and left to incubate overnight at 4°C. The following primary antibodies were used: IV58B6 mouse anti-glycogen antibody (1:500, Gentry Lab), rabbit anti-IBA-1 (1:500, Sigma, SAB5701363), goat anti-IBA-1 (1:500, Abcam, ab17884), and rabbit anti-pH2AX (1:500, Proteintech, 83307-2-RR). Sections were then washed in PBS for 5 x 5 minutes at RT, then incubated in 150µl with the appropriate Alexa Fluor conjugated secondary antibodies (all from Invitrogen and used at 1:1000 dilution): Donkey anti-goat AF-488 (A-11055), donkey anti-rabbit AF-488 (A-21206), donkey anti-rabbit AF-555 (A-31572), donkey anti-mouse AF-555 (A-31570), at RT in the dark for 2 hours. Following this, sections were incubated for 20 minutes at RT in the dark with DAPI (Thermo Scientific, 62248) at 1:1000 dilution in PBS. Slides were finally washed 5 x 5 minutes in PBS before coverslips were mounted with Fluoromount-G™ Mounting Medium (Invitrogen, 00-4958-02) and left to dry overnight in the dark before imaging.

### Image Analysis

Slides were imaged on a Nikon Ti2 Widefield Microscope. For each mouse, one slide containing five coronal brain sections was analysed. Two images were acquired from each section, one containing the hippocampus and one from a non-hippocampal region, yielding a total of 10 images per mouse. Image quantification was performed independently for each field of view, and measurements from all 10 images were averaged to generate a single value for each mouse. Individual mice, rather than individual images, were treated as the biological replicates for statistical analyses.

### Proteomics sample preparation

APP^NL-G-F^ and hAPP microglia were lysed in 400µl lysis buffer and *in vitro* microglia were lysed in 100µl of lysis buffer (5% sodium dodecyl sulfate (SDS), 10mM tris(2-carboxyethyl)phosphine (TCEP), 50mM Triethylammonium bicarbonate (TEAB)) and shaken at RT for 5 min at 1000rpm, followed by boiling at 95°C for 5 min at 500rpm. Samples were then shaken again at RT for 5 min at 1000rpm before being sonicated for 15 cycles of 30 s on/30 s off with a BioRuptor (Diagenode). Benzonase was added to each sample and incubated at 37°C for 15 min to digest DNA. Samples were then alkylated with 20mM iodoacetamide for 1h at 22°C. Protein concentration was determined using EZQ protein quantitation kit (Invitrogen, R33200) as per manufacturer instructions. Protein isolation and clean up was performed using S-TRAP (Protifi, C002-MINIX) micro spin columns before digestion with trypsin at 1:25 ratio (enzyme:protein) for 2h at 47°C. Digested peptides were eluted from S-TRAP columns using 50mM ammonium bicarbonate, followed by 0.2% aqueous formic acid and 50% aqueous acetonitrile containing 0.2% formic acid. Eluted peptides were dried down overnight.

### Mass Spectrometry

Peptides were run on an Orbitrap Astral mass spectrometer (Thermo Scientific) coupled to a Vanquish Neo UHPLC system (Thermo Scientific) with LC buffers compromising of buffer A (0.1% formic acid) and buffer B (80% acetonitrile (VWR, 83640.290), 0.1% formic acid). The buffers were used to create a gradient (4-35% B) for a 60SPD (samples per day) run where the peptides were eluted from a PepMap RSLC C18 column (Thermo Scientific, PNES906) and RAW data was acquired in Data Independent Acquisition (DIA) mode. With the Astral in positive mode, the voltage was set to 2.0 kV. A scan cycle compromised a full MS scan with an m/z range of 380-980, resolution of 240,000, custom Automatic Gain Control (AGC) target of 500% and a maximum injection time (IT) of 5ms. MS scans were followed by MS/MS DIA scans of dynamic window widths with an overlap of 0m/z. DIA spectra were recorded with a scan range of 150-2000m/z, custom AGC target of 500% and a maximum IT of 3ms. Normalised collision energy was set to 25% with a default charge state set at 2. Data for MS scans were acquired in profile mode with MS/MS DIA scan events being acquired in centroid mode.

### Proteomics data Handling, processing and analysis

Proteomes were analysed using Spectronaut (Biognosys) version 17. Raw mass spec files were searched against a mouse database (Swissprot Trembl November 2023) with the following parameters: directDIA, false discovery rate set to 1%, protein N-terminal acetylation and methionine oxidation were set as variable modifications and carbamidomethylation of cysteine residues was selected as a fixed modification.

### Downstream proteomic analysis

Proteomic data analyses were performed in R (version 4.5.2) with RStudio. Protein copy number data was filtered for known protein contaminants. All numeric sample data was log2 transformed before performing pairwise differential abundance analysis using the limma package (version 3.66.0). The pairwise results were filtered for missing values to confirm that each protein was detected in samples from both groups, with thresholds set to ensure there was ≥75% coverage for both groups. Significance was determined using FDR adjusted q-value and log2fold change thresholds, where proteins that significantly increased or decreased in abundance have a q.value < 0.05 and log2fold change of > 1.5 or <-1.5, respectively. To perform gene ontology (GO) analysis, Uniprot protein accession IDs were mapped to ENTREZIDs using the Bioconductor org.Mm.eg.db package (version 3.22.0). All proteins that passed NA filtering for the specific comparison between genotype groups were set as the background “gene universe” reference. Using the ClusterProfiler package (version 4.18.4), GO over-representation analyses were separately performed using significantly increased and decreased protein data subsets. Volcano and dot plots were made in RStudio using ggplot2 (version 4.0.2) and ClusterProfiler packages, respectively.

### Statistical analysis and data visualisation

Data were analysed using GraphPad Prism (version 10.6.1) and is reported as mean ± SEM unless otherwise stated. Students T-test and one-way ANOVA with Tukey’s post-hoc test were used unless otherwise stated in appropriate figure legends. Statistical significance was set at *p<0.05*.

## Data Availability

Raw mass spectrometry files have been deposited at ProteomeXchange and will be publicly available as of the date of publication.

---

**Supplemental Figure 1.**
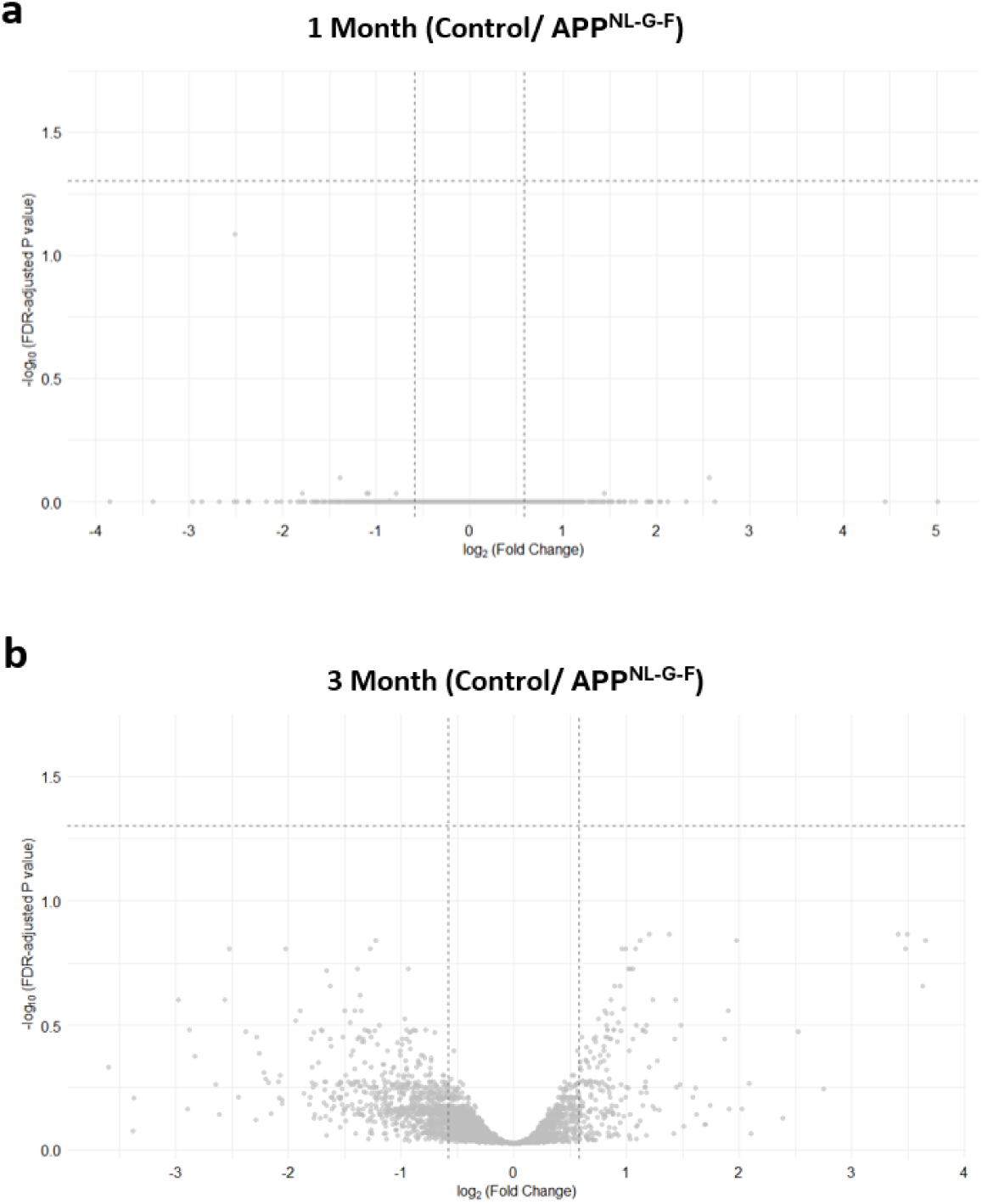
Mass spectrometry analysis shows no significant changes to microglia proteomes at early time points. **a)** Volcano plot of protein expression between 1-month APP^NL-G-F^ microglia and time-matched control. No proteins were significantly differentially expressed (FC > 1.5, FDR-adjusted P-value <0.05). **b)** Volcano plot of protein expression between 3-month APP^NL-G-F^ microglia and time-matched control. No proteins were significantly differentially expressed (FC > 1.5, FDR-adjusted P-value <0.05).

**Supplemental Figure 2.**
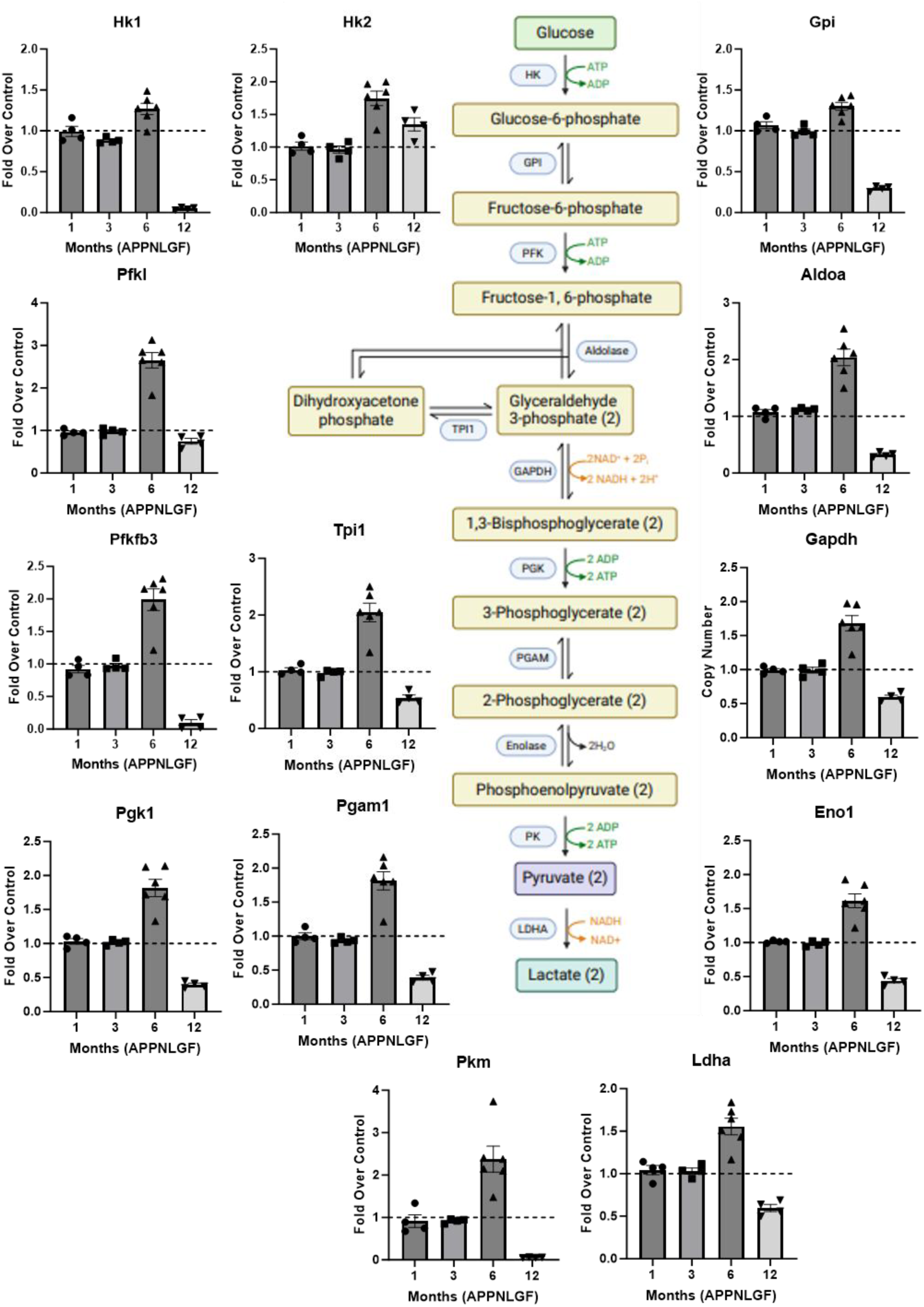
Mapping of glycolysis enzymes show dynamic metabolomic changes in APP^NL-G-F^ microglia. Copy number expression of all glycolysis enzymes in microglia from APP^NL-G-F^ mice presented as a fold-change over age-matched controls (dotted lines) ± S.E.M.

**Supplemental Figure 3.**
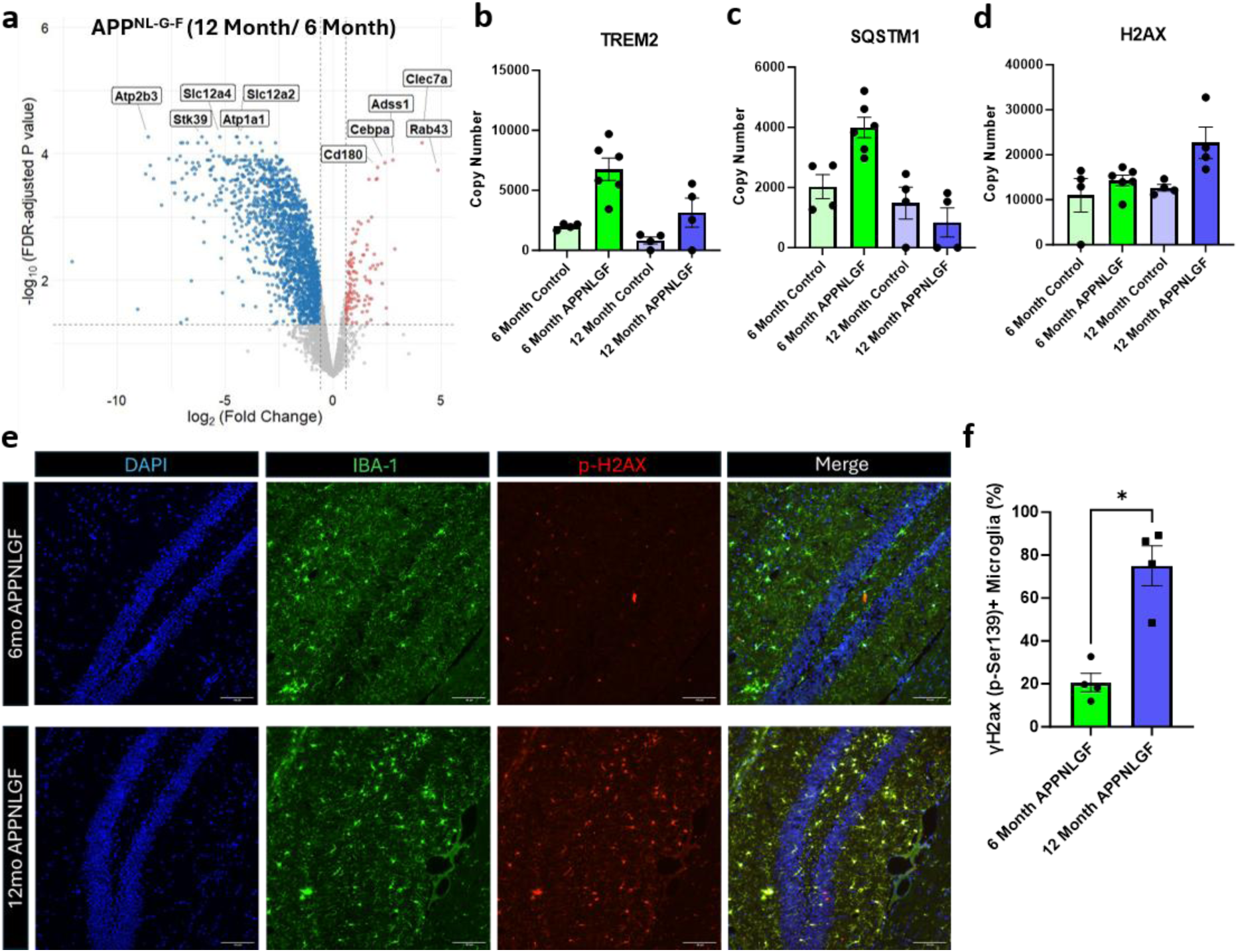
Late-stage disease microglia show signs of metabolic exhaustion and senescence. **a)** Volcano plot of differentially expressed proteins between 12-month (right) and 6-month (left) APP^NL-G-F^ microglia. Red dots = significantly upregulated proteins in 12-month APP^NL-G-F^ microglia (FC > 1.5, FDR-adjusted P-value <0.05), blue dots = significantly downregulated proteins in 12-month APP^NL-G-F^ microglia (FC > 1.5, FDR-adjusted P-value <0.05). **b)** Copy numbers of TREM2 in 6- and 12-month microglia from control and APP^NL-G-F^ mice ± S.E.M. N=3-4 per group. **c)** Copy numbers of Sequestosome 1 (SQSTM1; P62) in 6- and 12-month microglia from control and APP^NL-G-F^ mice ± S.E.M. N=3-4 per group**. d)** Copy numbers of H2AX in 6- and 12-month microglia from control and APP^NL-G-F^ mice ± S.E.M. N=3-4 per group**. e)** Representative images of brain sections from 6- and 12-month old APP^NL-G-F^ mice immunostained with DAPI (nucleus; blue), IBA-1 (green) and phospho12-month-old2AX (red). Scale bar = 100µm. **f)** Quantification of the percentage of γ-H2AX+ microglia (IBA-1+) in 6- and 12-month APP^NL-G-F^ brains ± S.E.M. N=4 per group.

**Supplemental Figure 4.**
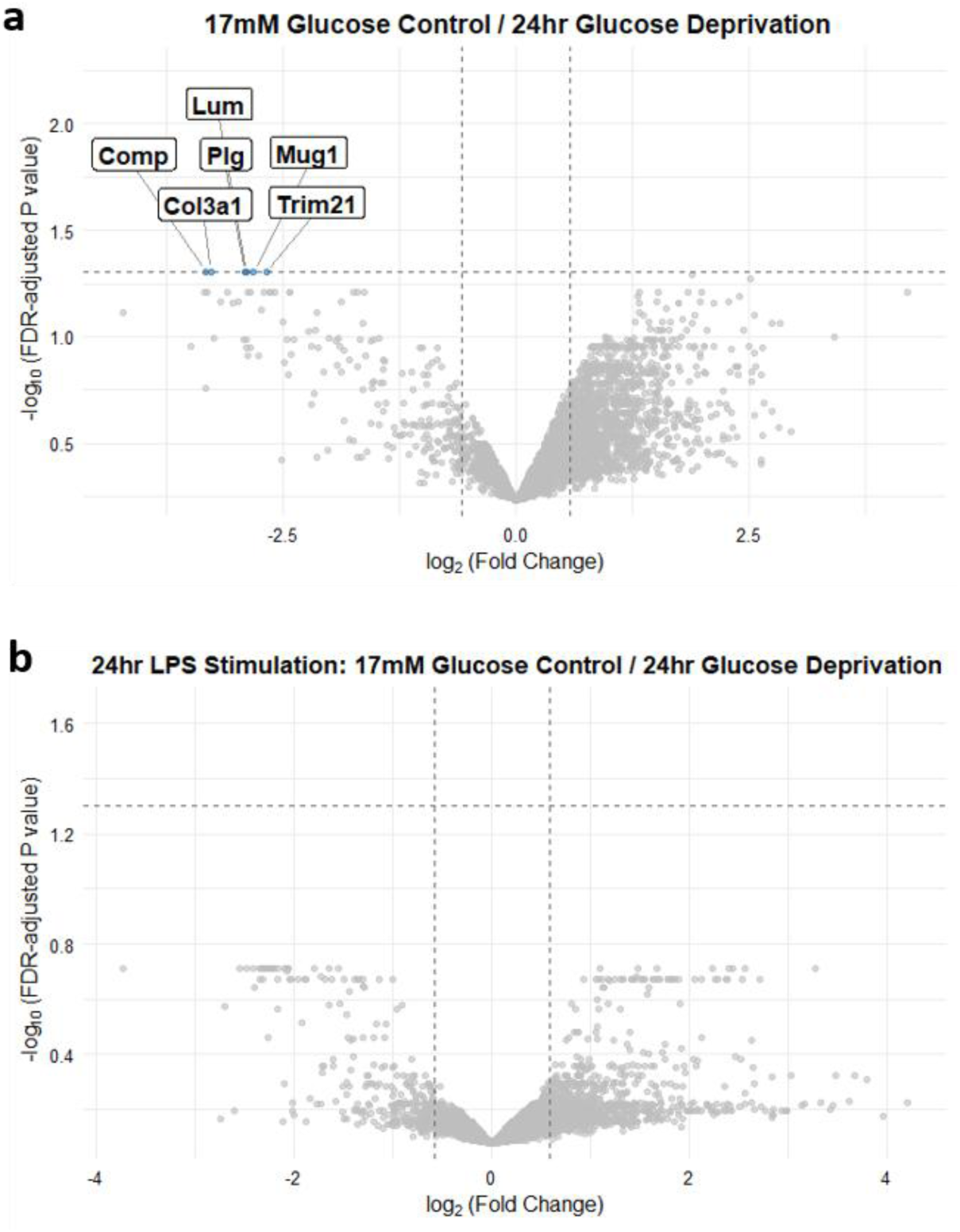
24-hour glucose deprivation does not impact microglia proteomes or responses to LPS. **a)** Volcano plot of protein expression between normal glucose (17mM) media (right) and glucose-depleted microglia (left). Blue dots = significantly downregulated proteins in glucose-deprived microglia. (FC > 1.5, FDR-adjusted P-value <0.05). **b)** Volcano plot of protein expression between LPS-treated microglia in normal glucose (17mM) media (right) and glucose-depleted media (left). No proteins were significantly differentially expressed (FC > 1.5, FDR-adjusted P-value <0.05).

**Supplemental Figure 5.**
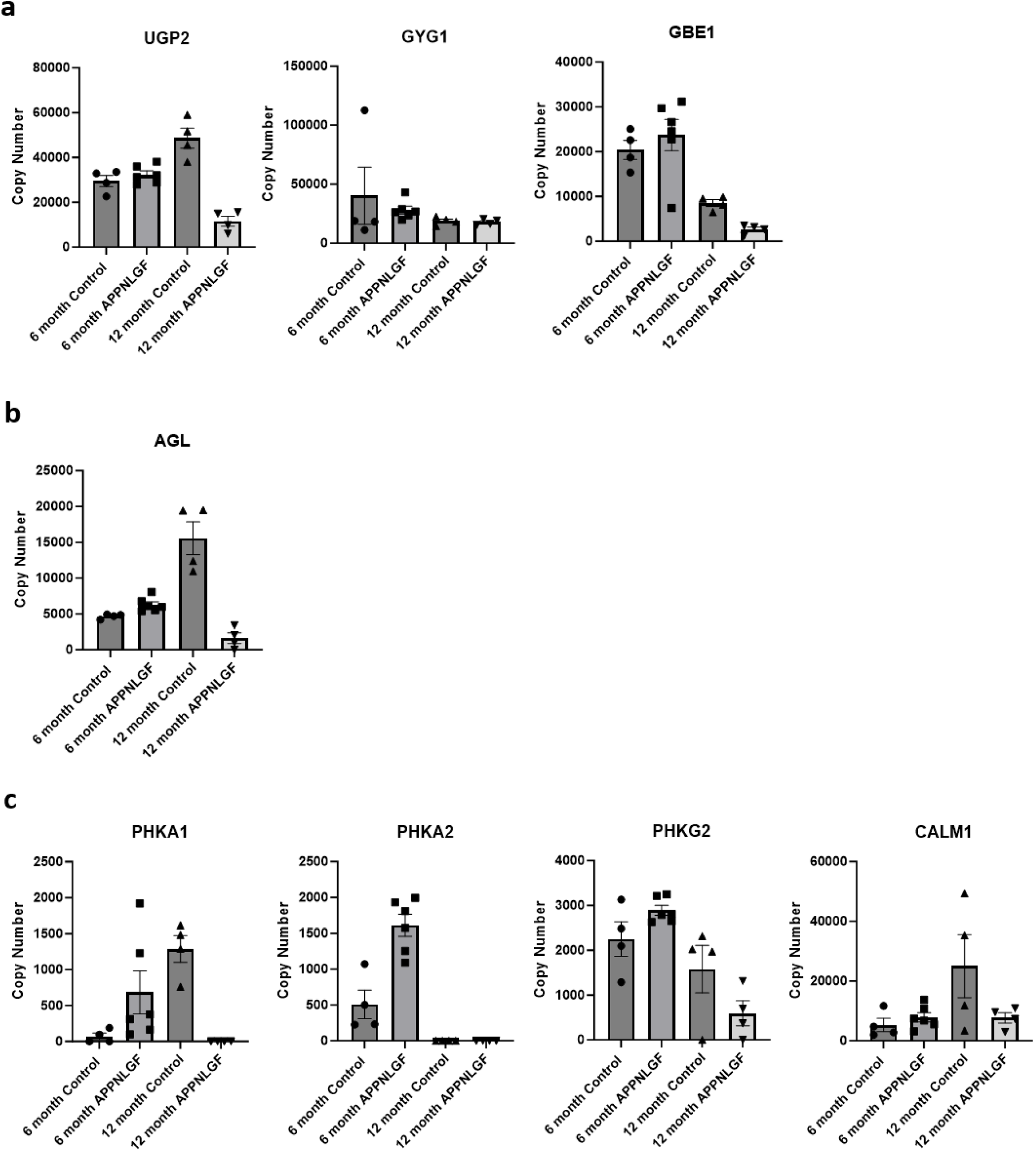
Microglia show age- and disease-associated changes to glycogen metabolism. **a)** Copy numbers of UDPG pyrophosphorylase (UGP2), Glycogenin (GYG1) and Glycogen Branching Enzyme (GBE1) in 6-month and 12-month old APP^NL-G-F^ and age-matched controls ± S.E.M. N=4-6. **b)** Copy numbers of Glycogen Debranching Enzyme (AGL) in 6-month and 12-month old APP^NL-G-F^ and age-matched controls ± S.E.M. N=4-6. **c)** Copy numbers of detected Phosphorylase Kinase subunits PHKA1, PHKA2, PHKG2 and CALM1 in 6-month and 12-month old APP^NL-G-F^ and age-matched controls ± S.E.M. N=4-6.

**Supplemental Figure 6.**
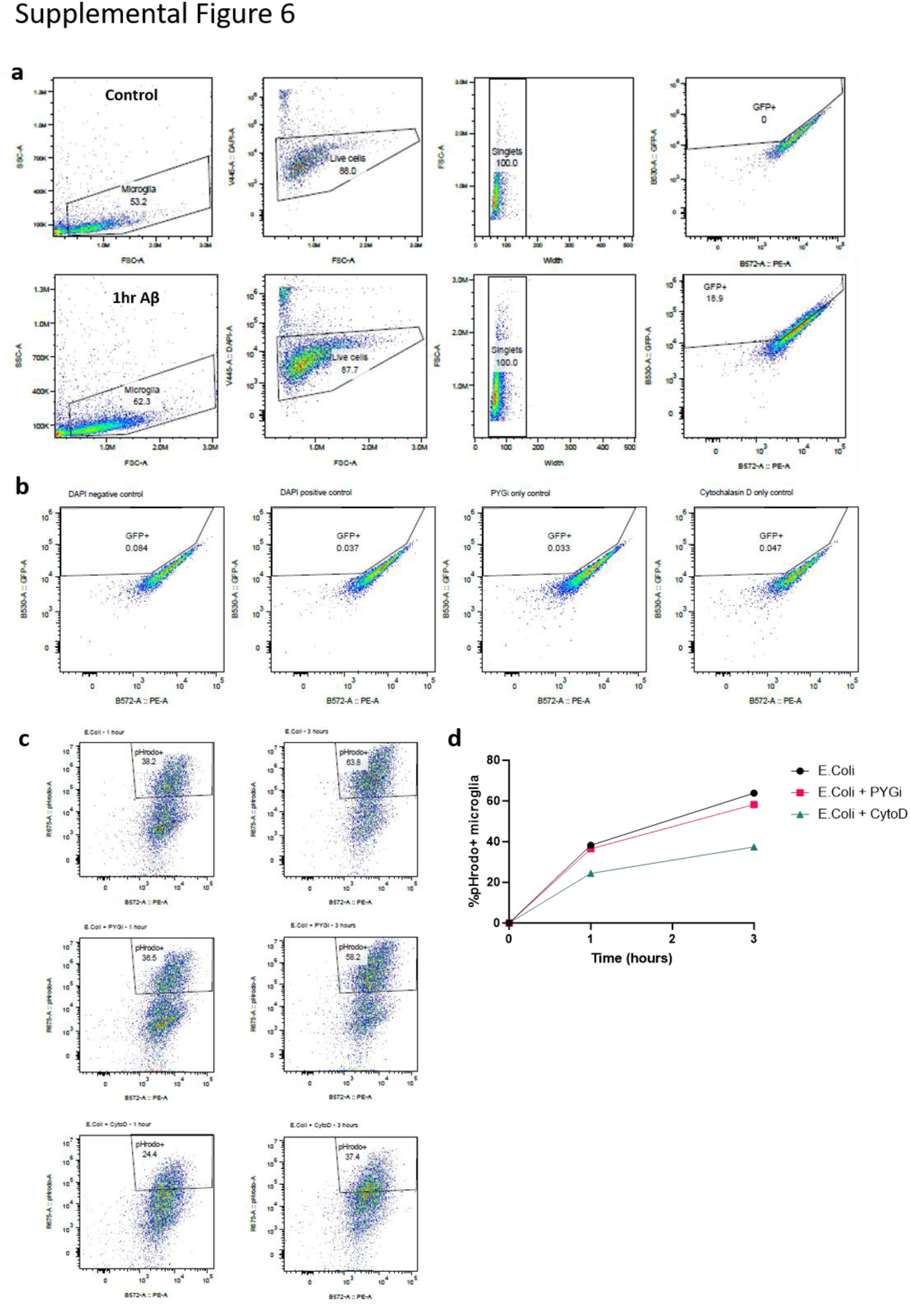
PYG inhibition does not reduce phagocytosis of E.coli particles. **a)** Flow cytometry gating strategy for phagocytosis assays. Top panel: gating strategies were performed on controls to firstly gate for cells, then viability using DAPI staining, followed by confirmation of singlets and then GFP expression. Bottom panel: example of gating strategy used on 1hr Aβ microglia to show shift in cell populations into GFP+ areas. **b)** Representative flow cytometry plots confirming no increases in GFP autofluorescence with inhibitors alone. Left plot: DAPI negative control; second plot: DAPI positive control; third plot: PYGi only control; fourth plot: Cyto-D only control. **c)** Representative flow cytometry plots to show pHrodo+ E. coli particle uptake in primary mouse microglia after 1 (left) and 3 hours (right). Top row: E. coli alone; middle row: E. coli + PYG inhibitor; bottom row: E. coli + Cyto-D. **d)** Quantification of pHrodo-E. coli+ microglia over time from flow cytometry analysis. N=1.

**Supplemental Figure 7.**
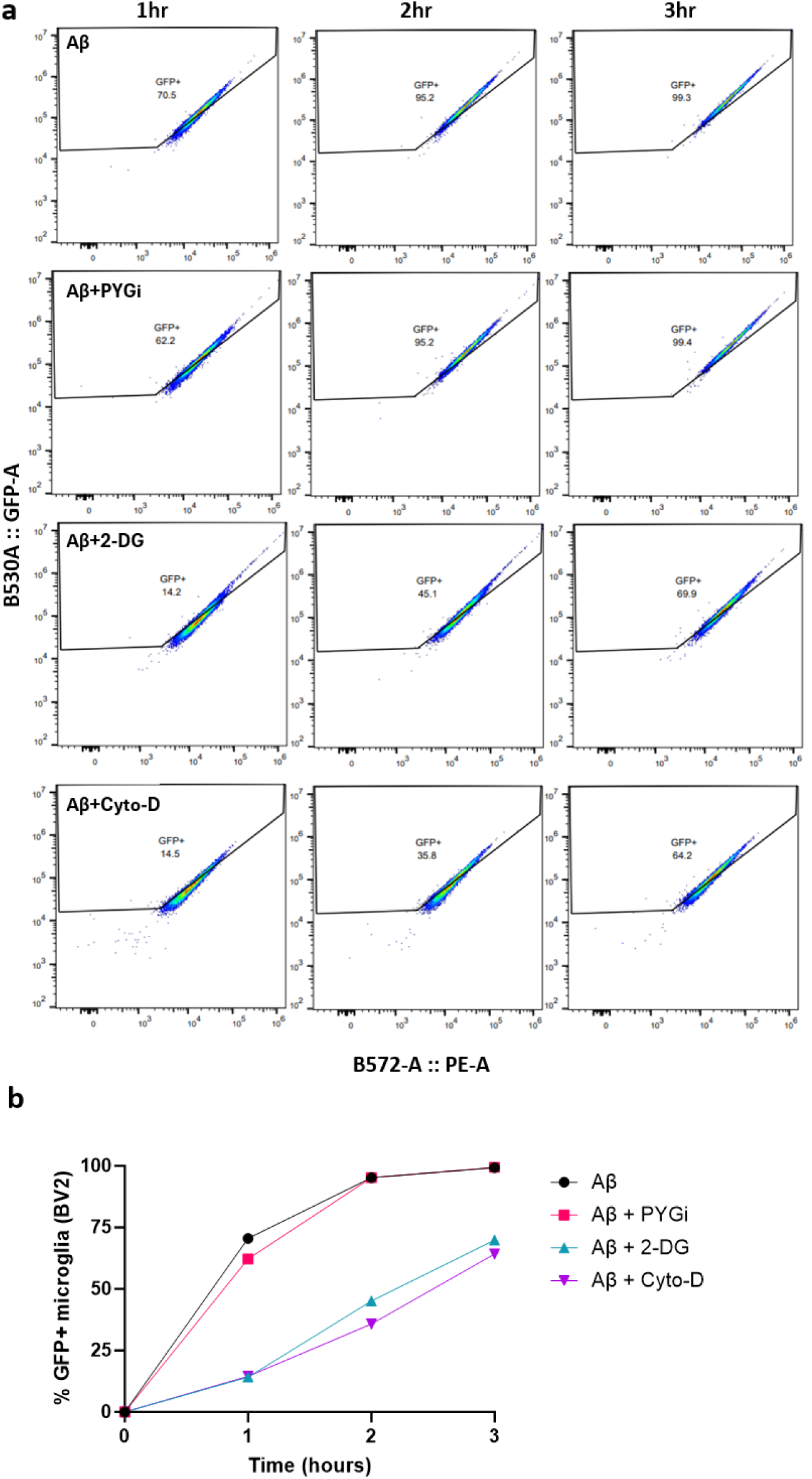
Glycogen phosphorylase inhibition does not impact Aβ uptake in BV2 cells. **a)** Flow cytometry plots of GFP+ microglia after 1, 2 and 3 hours of exposure to GFP-Aβ protein aggregates. BV2s were given GFP-Aβ aggregates alone (top row), with glycogen phosphorylase inhibitor (PYGi; second row), with 2-deoxyglycose (2-DG, third row), or with phagocytosis inhibitor Cytochalasin-D (Cyto-D; bottom row). **b)** Quantification of GFP+ microglia over time from flow cytometry analysis of GFP-Aβ aggregate uptake.

## Notes

### Competing Interest Statement

The authors have declared no competing interest.

### Funding

- Alzheimer’s Research UK: ARUK-RADF2022B-004

